# Photoacoustic imaging assessment of acute kidney injury associated with experimental necrotizing enterocolitis

**DOI:** 10.1101/2025.09.30.679399

**Authors:** Anchala Guglani, Angadh Singh, Maryssa A. Ellison, Nildris Cruz-Diaz, Parvesh M. Garg, Jared A. Weis, Victoria G. Weis

## Abstract

**Background:** Necrotizing enterocolitis (NEC) is a devastating gastrointestinal condition increasingly recognized to cause systemic complications, including acute kidney injury (AKI). However, the underlying mechanisms and variations in renal impairment remain largely unknown, and there is a lack of non-invasive imaging methods for detecting early kidney injury.

**Methods:** We evaluated kidney injury through a multimodal approach, using a neonatal rat model of experimental NEC. Plasma and renal tissue were analyzed for pro-inflammatory cytokines (TNF-α, IL-6, IL-1β) by ELISA. Histological analysis and immunofluorescence staining were used to identify renal abnormalities and expressions of KIM-1 and NGAL. Photoacoustic imaging (PAI) was used to assess renal oxygenation and total hemoglobin as functional imaging biomarkers of kidney injury.

**Results:** NEC pups showed significant elevations of systemic and renal cytokines, as well as increased expression of KIM-1, NGAL and CD31 in proximal tubules of the kidney. Histological examination further confirms renal injury in the NEC cohort. PAI demonstrated reduction in renal oxygen saturation, offering non-invasive assessment of physiological kidney damage.

**Conclusion:** This study characterizes AKI associated with NEC as an inflammatory and hypoxic response in the neonatal rat NEC model and introduces PAI as a promising imaging modality for non-invasive evaluation of renal injury in NEC.

## 1. Introduction

Necrotizing enterocolitis (NEC) is one of the most devastating gastrointestinal conditions, predominantly impacting preterm and extremely low birth weight infants causing significant morbidity and mortality in neonatal intensive care units around the world(1–3). NEC affects approximately 6-10% of this vulnerable population. The prevalence and severity of NEC continues to rise despite advancements in neonatal intensive care(1, 4, 5). The disease is characterized by rapid progression from intestinal inflammation and disruption of the epithelial barrier to extensive tissue necrosis that leads to intestinal perforation followed by sepsis-induced multisystem organ failure. While the mechanisms driving the onset and progression of NEC are still being uncovered, current evidence suggests that the pathogenesis of the disease is likely caused by a combination of bacterial infection and hyperinflammatory responses that are aggravated by multiple factors, such as prematurity, formula feeding, hypoxia, and impaired intestinal barrier function(1, 6, 7). The impact of early-stage damage on the bowel can eventually lead to complex systemic effects, with both diagnosis and intervention particularly challenging in premature infants(8, 9).

The pathophysiology of NEC has been mainly attributed to the gastrointestinal tract. However, recent research suggests that NEC-associated inflammation potentially induces injury in organs beyond the intestine, including the kidney, liver, and brain(10–13). It has been clinically and experimentally documented that NEC can lead to multi-organ dysfunction syndrome due to increased inflammatory mediators, breakdown of the intestinal barrier, and subsequent bacterial translocation(14). Among extraintestinal organs, the kidney is highly susceptible to injury due to its connected microvasculature, high metabolic demand, and sensitivity to changes in oxygenation and perfusion. Acute kidney injury (AKI) is one of the most significant extraintestinal complications of NEC. AKI has been reported in up to 60% of neonates with NEC and is associated with increased mortality, prolonged hospitalization, and adverse neurodevelopmental consequences(13, 15). Both short-term complications and long-term renal impairment are associated with the degree of kidney involvement following NEC(16). NEC and AKI have complex mechanisms, involving systemic hypoperfusion, inflammatory cytokines, oxidative stress, and exposure to nephrotoxic agents, all factors that collectively cause tubular epithelial damage and renal microvascular injury. Despite frequent observations of renal dysfunction in infants with NEC, the underlying mechanisms, timing, and variations in renal impairment remain largely unknown. As a consequence, there are no direct functional or molecular measures linking NEC to in vivo kidney alterations(17). Further aggravating the problem, clinical biomarkers for AKI detection such as serum creatinine are not adequate for diagnosis in neonates(18, 19). Thus, there is growing interest in early biomarkers such as kidney injury molecule-1 (KIM-1), which is specifically upregulated in hypoxic and inflammatory conditions. Studies demonstrate that KIM-1 and other biomarkers, such as neutrophil gelatinase-associated lipocalin (NGAL), are more sensitive than conventional measures for early detection of AKI(19–22). Although these biomarkers are molecular indicators of damage, they are unable to identify dynamic and causative shifts in renal perfusion and oxygenation that occur prior to structural damage.

Advancing our current understanding of NEC-associated kidney damage requires development of new imaging methods that can measure these functional characteristics in vivo. Conventional imaging methods typically available for the assessment of NEC and related injuries are X-rays and ultrasound but possess limited degrees of specificity and sensitivity for diagnosing early intestinal or extraintestinal damage(23, 24). In neonatal rat models, magnetic resonance imaging (MRI) has shown promise in assessing intestinal inflammation using variables like T2 relaxation time and the apparent diffusion coefficient (ADC), which has a significant correlation with histological damage. However, use of MRI in neonatal intensive care facilities is limited by its high cost and restricted accessibility due to patient transportation issues(25). To address these challenges, non-invasive functional imaging approaches that analyze oxygenation, hemodynamics, and microvascular integrity are required. Recent advances in the field of noninvasive imaging have introduced photoacoustic imaging (PAI). PAI is an innovative approach to non-invasively measuring tissue oxygenation and hemodynamics that combines optical contrast with ultrasonic detection. Through the use of this emerging imaging technology, tissue oxygen saturation and perfusion can be measured in real-time and at high resolution using hemoglobin optical absorption properties(26). In recent work, we utilized PAI in preclinical NEC studies and demonstrated that PAI could sensitively detect reductions in intestinal oxygen saturation at both early and late stages of NEC disease in a neonatal rat experimental model of NEC(27). These findings were confirmed by intestinal transit assay and histology, validating PAI as a promising tool for early detection of NEC. Importantly, this provides evidence that real-time, quantitative functional measures of tissue oxygenation via PAI directly correlate with tissue-level disease severity(27).

Despite advances in PAI, few studies have applied PAI to evaluate microvasculature and oxygen saturation as a measure of renal health and function. Previous studies have demonstrated that PAI detects substantial reductions in renal oxygen saturation which were associated with histological indications of tubular injury and subsequent functional impairment in preclinical ischemia reperfusion models(28). Further advances in label-free dual-modal photoacoustic/ultrasound localization imaging have allowed three-dimensional mapping of vascular density, relative blood volume, and oxygenation in experimental AKI, with marked reductions found in these parameters(29). In general, these studies illustrate the promise of PAI assessment of renal oxygen saturation and hemodynamics as an analytical tool for characterizing renal injury(28, 29). However, no studies have been reported for the use of PAI in NEC-associated AKI despite the pathophysiological overlaps between hypoxia, inflammation, and microvascular dysfunction. Extending PAI assessment to kidney injury associated with NEC has the potential for novel and translational advancement in a vulnerable population where early non-invasive detection can significantly improve long-term health outcomes. Considering these developments and our previous studies on PAI in experimental NEC(26, 27), the rationale for this study is to characterize NEC-associated acute kidney injury using multimodal methods during the course of NEC. The objectives of our study are to utilize photoacoustic imaging coupled with quantitative biochemical assays, histopathological, and immunofluorescence analyses in rat neonatal NEC models to provide a comprehensive evaluation of NEC-associated AKI. We hypothesized that NEC-induced systemic inflammation and tissue hypoxia will result in kidney dysfunction and injury that can be detected by PAI and validated by elevated plasma and tissue cytokine levels and histological markers of tubular damage. Through this integrated approach, bridging both basic and translational science, our study seeks to provide a foundation for improvement in the diagnosis and treatment of preterm infants with NEC-associated AKI.

## 2. Materials and Methods

### 2.1 Animals and Experimental Design

Animal experiments were performed according to protocols approved by the Institutional Animal Care and Use Committee (IACUC) of Wake Forest University. Timed-pregnant Sprague-Dawley dams were procured from Charles River Laboratories (Wilmington, MA) for this study. Dams were monitored 36 hours before delivery was expected. After birth, pups were randomly divided into experimental NEC or breastfed (BF) control cohorts. In the NEC cohort, pups were isolated from dams within 6 hours of birth and housed in a clinical grade neonatal incubator adapted for animal studies to maintain consistent temperatures (32°C) and humidity (>50%) throughout the study period, placed in a dedicated room within the vivarium space. NEC pups were subjected to the NEC experimental model for 4-days via hypertonic formula feeding (four times per day), full-body hypoxic stress (5% O2, three times per day), and orally dosed lipopolysaccharide to mimic bacterial stress (4 μg/g, three times within 30 hours of birth) for NEC induction, as previously described(30). BF pups remained with their dams in the same isolated rooms as NEC pups for the duration of studies. BF pups were removed from dams briefly for imaging procedures and returned to dams following anesthesia recovery. All pups were observed for clinical signs, including weight changes and feeding intolerance, until euthanasia performed on the fourth day post-birth. Body weight (BW) and Clinical Sickness Score (CSS) were measured at birth and every 24 hours. The Clinical Sickness Score combines four criteria: appearance, touch response, natural activity, and body color as previously described(30, 31).

### 2.2 Ultrasound and photoacoustic imaging

A Vevo F2 LAZR-X high-resolution ultrasound and photoacoustic imaging system (FUJIFILM VisualSonics, Inc., Toronto, ON, Canada) was used to collect high frequency anatomical US and PAI images. Anesthesia was induced in rat pups with the use of an anesthesia vaporizer system using 2% isoflurane inhalation in medical air. After induction, anesthesia was maintained at 1% isoflurane in medical air throughout all imaging studies. Animals were positioned supine on a heated platform with continuous physiological monitoring and imaging was acquired as previously described(27). Ultrasound gel was used to provide acoustic coupling with the abdomen of rat pups. Imaging acquisition methods and parameters were held constant among all experimental animals used in this study. Renal oxygenation and blood flow were measured in 4-day old rat pups with PAI imaging acquired at spectral excitation wavelengths of 750 nm and 850 nm in Vevo ‘OxyHemo’ mode with a two-dimensional sagittal imaging plane centered at the kidney midplane. Assessment of renal tissue oxygenation (sO2) and total hemoglobin (HbT) is based on automatic calculation of oxy- and deoxyhemoglobin concentrations within VevoLab postprocessing analysis software (FUJIFILM VisualSonics, Inc., Toronto, ON, Canada) using OxyZated and HemoMeaZure analysis tools, respectively. Mean sO2 and HbT measurements were quantified by manually drawing a region-of-interest (ROI) over the kidney region for each animal based on the anatomical ultrasound B-mode images. Similar to our previous study(27), NEC pups were imaged a minimum of 6 hours after the last total body hypoxia session that was used to induce NEC disease to avoid interference with sO2 measures by allowing for systemic equilibration of blood oxygenation.

### 2.3 Histological and Immunofluorescence Analysis

At the 4-day timepoint, kidney samples were harvested from healthy and NEC cohorts and processed for histological and immunofluorescence analysis as described in our previous study(30). Hematoxylin and eosin (H&E) staining was carried out on 4% paraformaldehyde-fixed, paraffin-embedded 5 μm sections and imaged at 20× magnification using an EVOS FL Auto 2 Cell Imaging System (Invitrogen, CA). H&E-stained sections were examined to identify morphological alterations including tubular dilatation, interstitial edema, inflammatory infiltrate, brush border loss, and tubular necrosis(32). Immunofluorescence (IF) analysis was performed on kidney sections after deparaffinization, rehydration, and antigen retrieval using Antigen Retrieval Buffer (Abcam, MA) as previously described(30). After blocking with protein block serum-free (Dako, Denmark), sections were incubated with primary antibodies Goat anti-KIM (Cat No. AF3689, R&D Systems, Minneapolis, MN), Rabbit anti-NGAL (Cat No. 39141T, PA5-79590; Thermo Fisher Scientific), and Goat anti-CD31 (Cat No. AF3628, R&D Systems, Minneapolis, MN) overnight at 4°C. Secondary antibodies conjugated for immunofluorescence with Alexa Fluor 488 and 549 (Jackson ImmunoResearch, PA) were used and then sections were incubated with 4′,6-diamidino-2-phenylindole (DAPI). Fluorescence images were acquired using a fluorescence microscope (EVOS FL Auto 2, Invitrogen). Image quantification was performed using ImageJ software. For IF images, mean fluorescence intensity was quantified from specified regions of interest following background subtraction and averaged across images for each sample. H&E images were evaluated qualitatively, and representative areas of morphological changes were indicated with arrows without further quantitative analysis.

### 2.4 Assessment of Biochemical Markers in Plasma and Tissue Samples

Rat pups were euthanized by rapid decapitation at end point (day 4) according to protocols approved by the IACUC. Blood samples were collected immediately into EDTA-coated tubes and centrifuged at 10,000 × g for 10 min at 4°C. Plasma samples were stored in clean tubes for further studies. Following the manufacturer’s instructions, plasma samples were analyzed for inflammatory cytokines, including tumor necrosis factor-α (TNF-α), interleukin-6 (IL-6), and interleukin-1β (IL-1β) using enzyme-linked immunosorbent assay (ELISA) kits (CSB-E11987r, CSB-E04640r CUSABIO, Houston, USA, and RLB00-1 Quantikine, R&D Systems, MN, US). Renal inflammation was quantified for the same cytokines in homogenized kidney tissue samples using ELISA. Assays were performed in duplicates to ensure accuracy, using high specificity and sensitivity ELISA kits. Kidney function biomarkers including blood urea nitrogen (BUN) and creatinine levels were measured from plasma samples using commercial assay kits, following the manufacturer’s instructions and performed by the Wake Forest University School of Medicine Biomarker Analytical Core.

### 2.5 Statistical Analysis

The change in body weight and clinical sickness score data was analyzed by two-way ANOVA. PAI measurements of mean kidney tissue oxygenation and mean total hemoglobin were compared between the BF and NEC cohorts using an unpaired t-test. ELISA measures of inflammatory cytokines were compared using unpaired t test with Welch’s correction. All analyses were conducted using GraphPad Prism software with a *p* value less than 0.05 considered as statistically significant.

## 3. Results

### 3.1 Confirmation of NEC Induction and Intestinal Damage

The experimental NEC animal model successfully induced NEC in neonatal rats using formula feeding, LPS administration, and hypoxic stress within 6 hours of birth. Previous studies suggest that intestinal damage begins within 24 hours of NEC induction, whereas overall mortality limits the study period to 96 hours(33). In our experimental cohort, BW and CSS in NEC and BF animals throughout study duration were monitored. CSS was assessed daily using criteria such as color, response to touch, appearance, and activity level (0–3 per domain), with higher scores signifying worsening clinical symptoms(34). Within 24 hours of exposure to NEC conditions, NEC pups showed decreased BW and elevated CSS, with these symptoms worsening over time until the end point of the study (Figure 1). BF pups gained weight and did not exhibit any significant clinical sickness signs from birth to end point. These results are consistent with our previous study which demonstrated that intestinal injury is corelated with weight loss and increased CSS(30).

**Figure 1.**
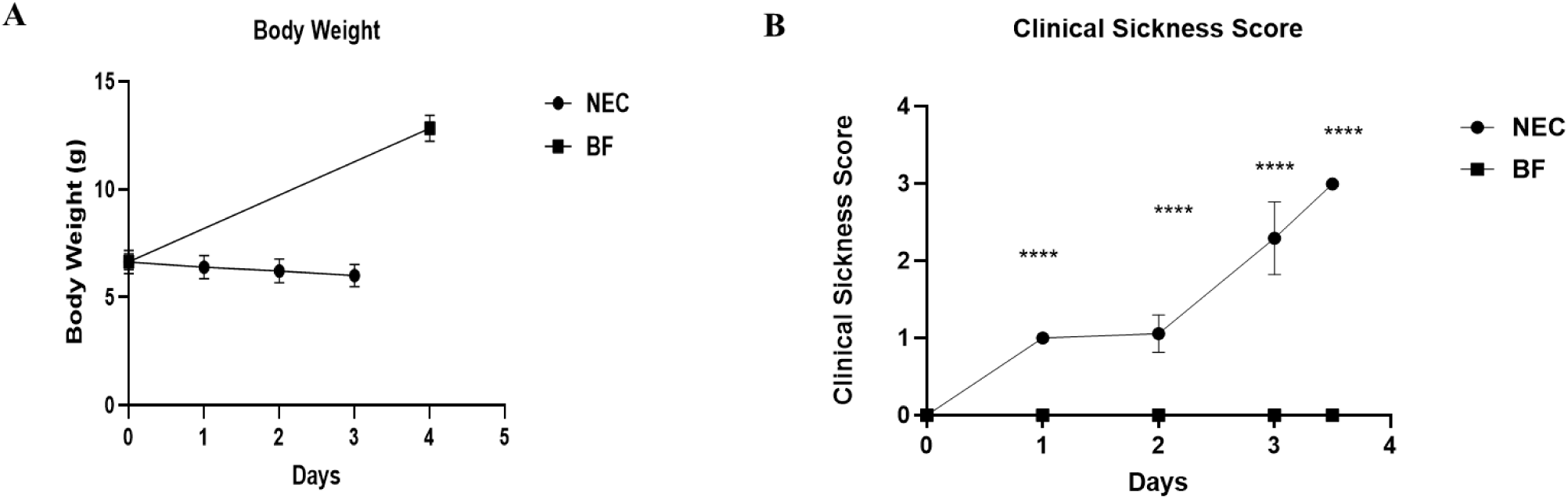
Assessments of body weight and clinical sickness score. **(A)** Body weight of BF and NEC cohort (mean ± SEM); NEC animals showed progressive weight loss relative to BF. NEC pups were weighed daily from day 0 through day 3 and BF pups were weighed on day 0 and day 4, as they remained with the dam and could not be handled on intermediate days **(B)** Clinical sickness scores on a 0 - 3 scale in BF and NEC cohorts from day 0 to day 4 (mean ± SEM). NEC pups exhibited significantly higher sickness scores compared to BF **(**** *p* < 0.0001)**.

### 3.2 Plasma and Renal Inflammation with Renal Function Measures in NEC Pups

Plasma cytokine levels were quantified by ELISA to determine whether NEC induction was linked to systemic inflammation. The plasma samples of NEC pups showed significantly higher concentrations of TNF-α, IL-6, and IL-1β in comparison to the BF pups plasma samples (\**p* < 0.05) (Figure 2A). Additionally, TNF-α, IL-6, and IL-1β concentrations were significantly greater in kidney homogenates from NEC pups than those of BF controls (\*\**p* < 0.01) (Figure 2B), suggesting that local renal inflammatory responses mimicked systemic inflammation. Renal function biomarkers, including blood urea nitrogen (BUN) and creatinine, were assessed in plasma samples. There were no significant differences observed between BF and NEC cohorts on day 4 (Figure 3). This finding aligns with the previously reported limitation of these renal function parameters in neonates for detecting early AKI(19).

**Figure 2.**
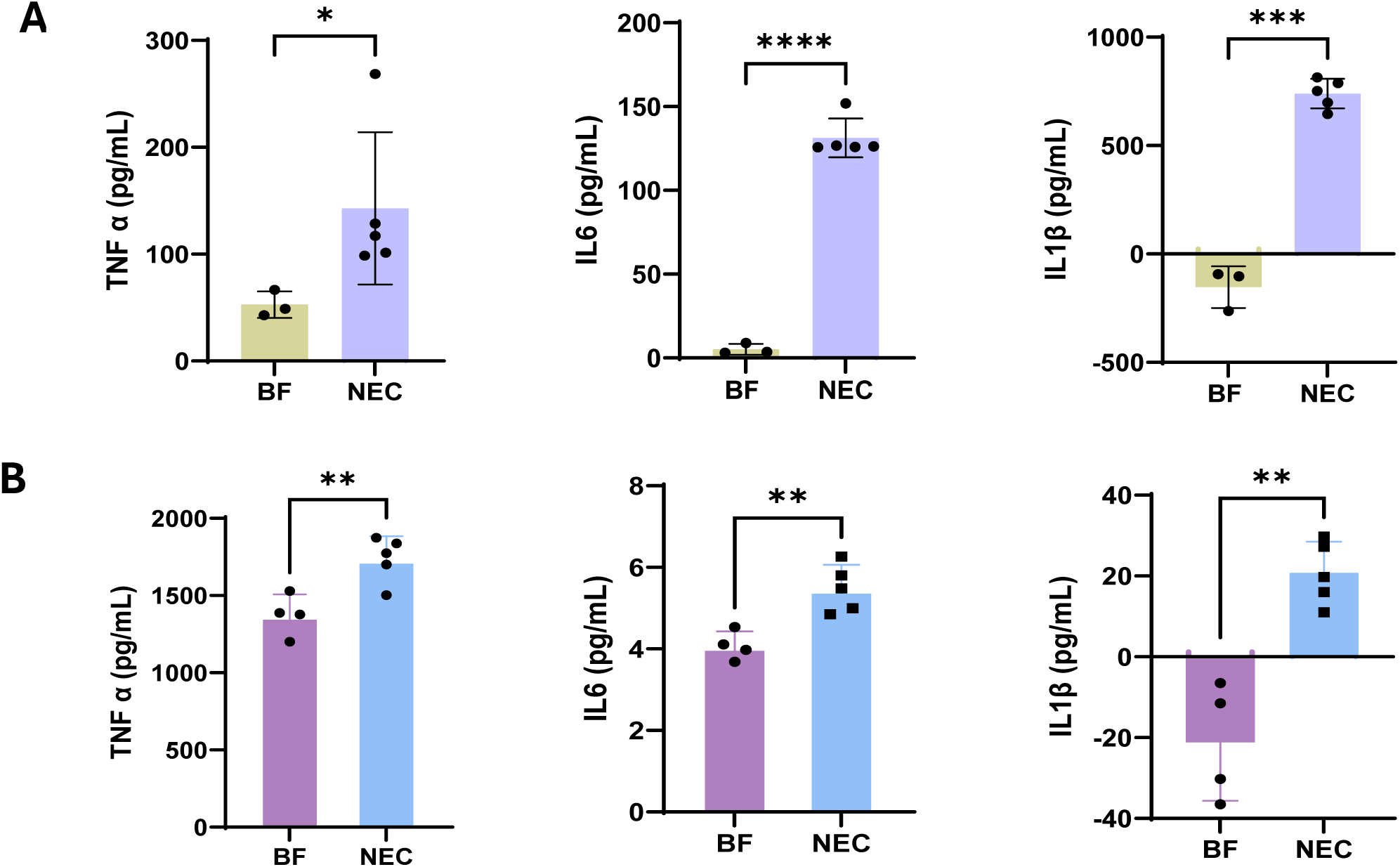
Plasma and kidney cytokine levels in BF and NEC cohort. Concentrations of TNF-α, IL-6, and IL-1 β were quantified by ELISA in plasma **(A)** and kidney tissue homogenates **(B)** from BF and NEC pups. Data are presented as Mean ± SEM. Statistical analysis was performed using an unpaired t test with Welch’s correction, with p values indicated on the graphs **(**** *p* < 0.0001, \*\*\**p* < 0.001, \*\**p* < 0.01, *p < 0.05).**

**Figure 3.**
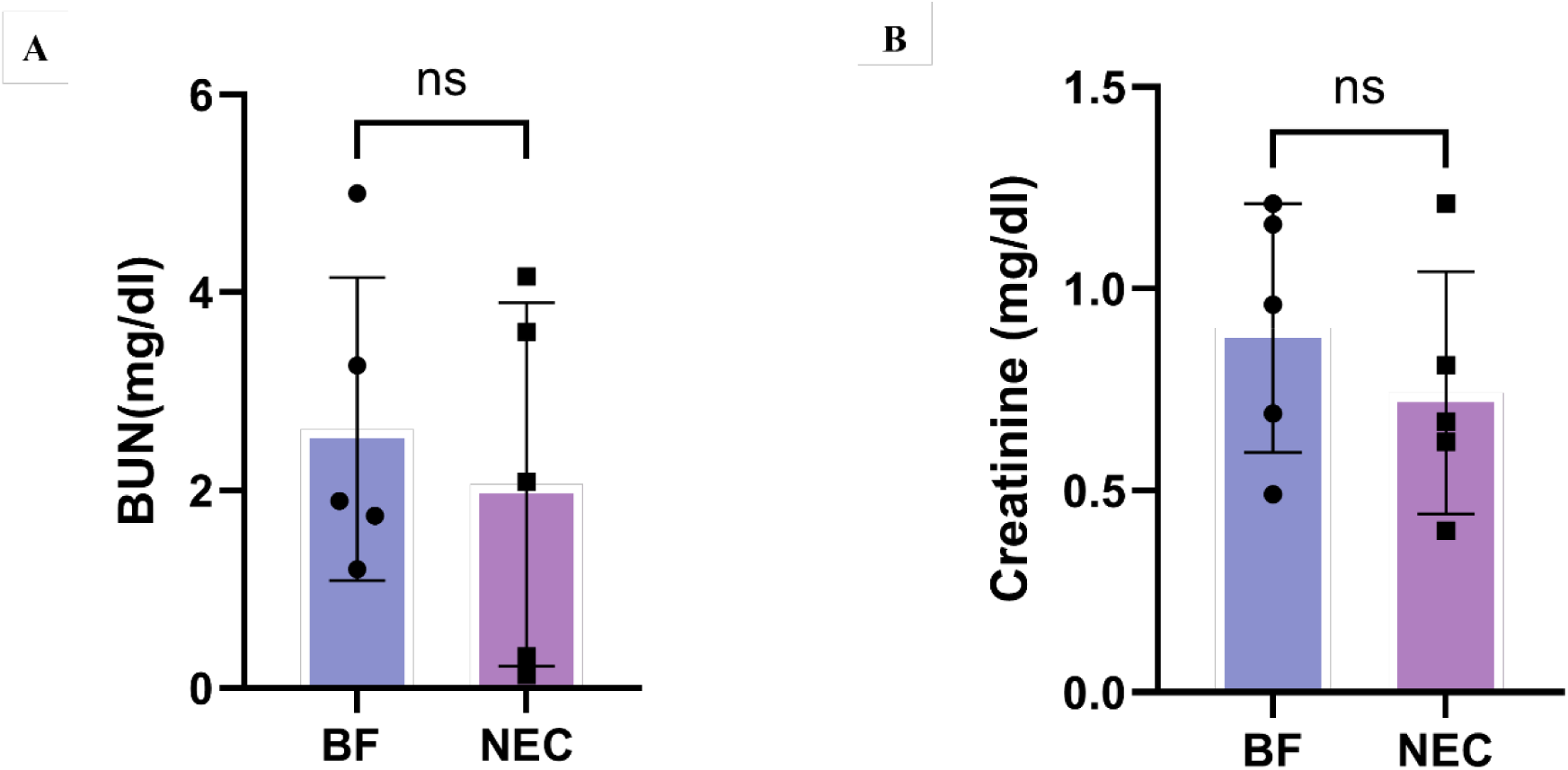
Plasma biochemical markers in BF and NEC cohort. **(A)** Plasma blood urea nitrogen (BUN) and **(B)** creatinine concentrations were measured in BF and NEC pups. Data are presented as mean ± SEM. Comparisons between groups were made using an unpaired t-test with Welch’s correction. ns = no statistically significant differences were observed among groups.

### 3.3 Histological Examinations of Kidney

To further confirm AKI associated with NEC, H&E staining of kidney tissue from BF and NEC pups were examined. Histological analysis of kidney tissue sections from NEC pups showed tubular dilation and interstitial edema as compared to BF kidney tissue (Figure 4). These characteristics are indicative of acute tubular damage and were not observed in BF. Acute tubular injury was confirmed by tubular dilation, cellular edema and inflammatory cell infiltration, similar to a previous study(32). In addition, IF staining was performed using KIM-1 and NGAL kidney injury biomarkers to characterize renal injury, along with CD31 staining to evaluate vascular endothelial cell expression and renal microvasculature architecture. IF images show that all markers demonstrate high intensity in kidney tissue from NEC pups (Figure 5 and Figure 6). KIM-1 expression was observed predominantly in proximal tubular epithelial cells and NGAL staining was markedly observed in the tubular compartment of NEC kidneys. These observations reflect ongoing tubular injury and early injury responses in NEC kidneys, while the BF control kidneys showed absence of these molecular biomarkers, reflecting intact tubular integrity. CD31 positive endothelial cells demonstrated significantly higher expression in kidney tissue from NEC pups compared to BF controls (Figure 6). CD31 staining was markedly observed in both glomerular capillaries and peritubular microvasculature of NEC kidneys, reflecting an upregulated endothelial response. In contrast, BF control kidneys showed lower CD31 expression with preserved vascular architecture. The presence of KIM-1 and NGAL in NEC tissue validates the histological results of tubular damage and indicates AKI. The above findings illustrate that NEC causes substantial endothelial cell activation and vascular remodeling alongside the renal epithelial damage evaluated at the structural and molecular levels.

**Figure 4.**
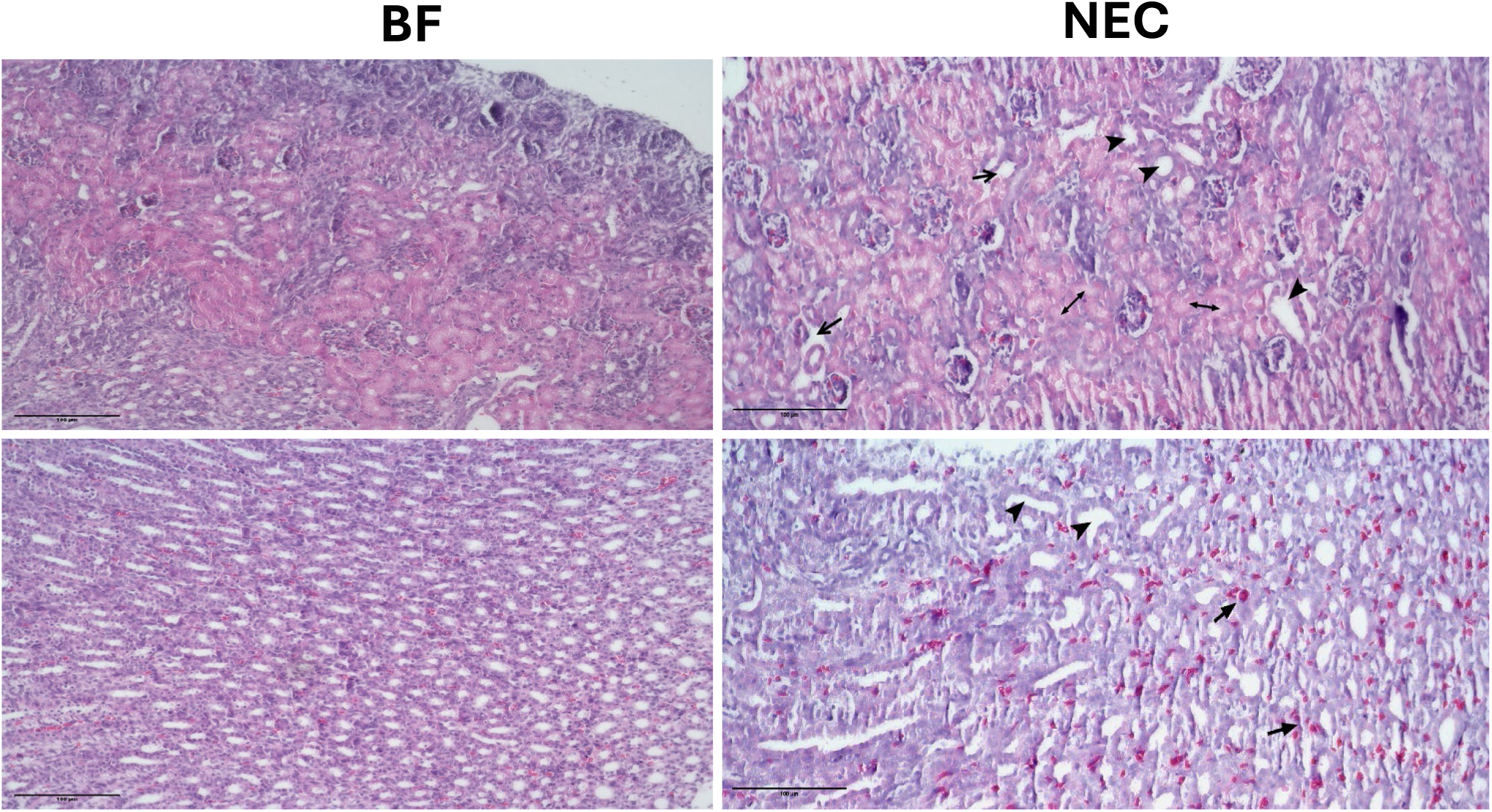
Histological abnormalities in the kidneys of BF and NEC pups. Representative images of H&E-stained kidney sections indicating tubular dilation, interstitial edema and tubular epithelial inflammation.

**Figure 5.**
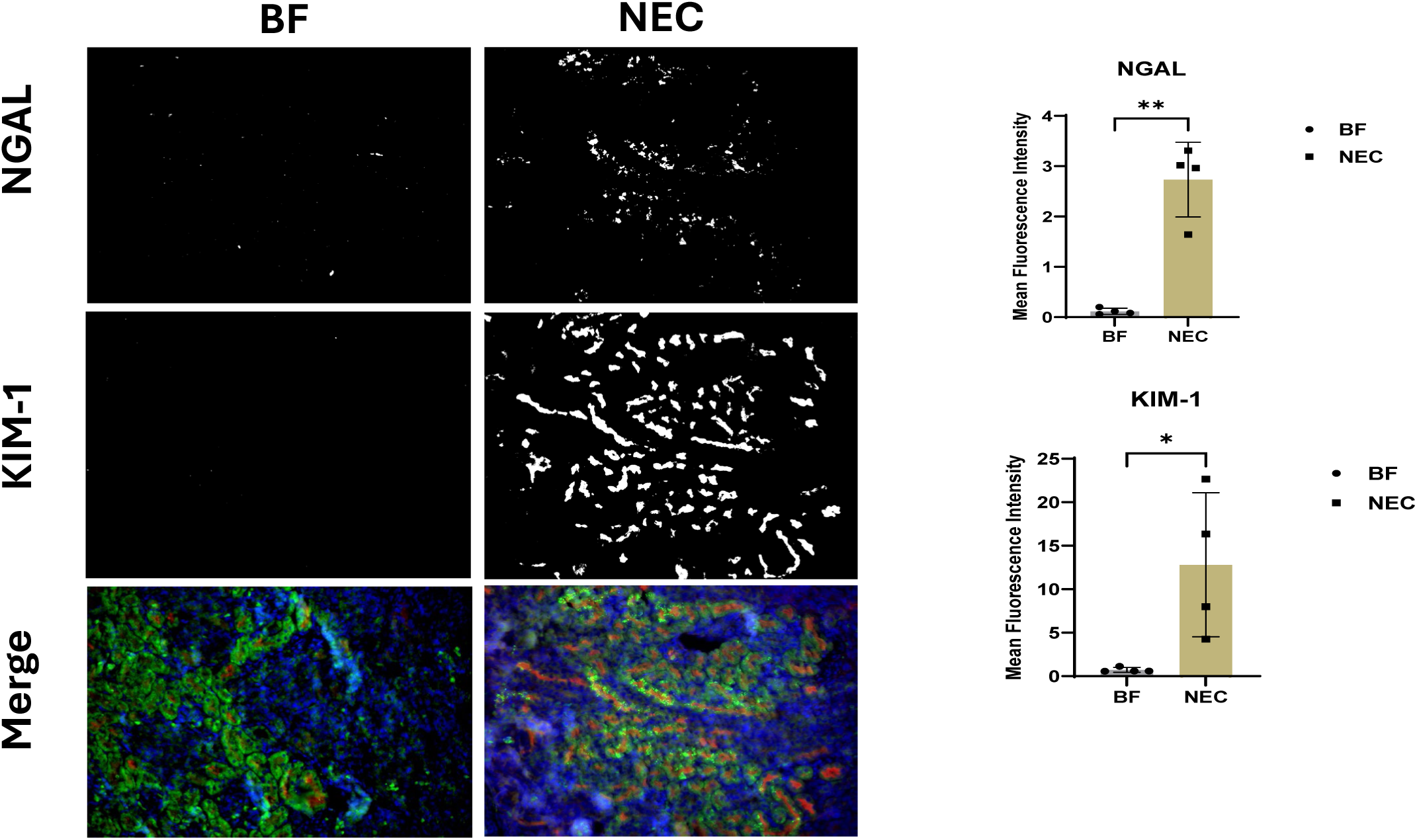
Expression of KIM-1 and NGAL in BF and NEC kidney tissue. **(Left)** Representative images of immunofluorescence staining of (**A**) NGAL and **(B)** KIM-1. BF kidney sections showing negative background staining for NGAL and KIM-1, NEC sections showing strong focal KIM-1 and NGAL expression. **(Right)** Quantitative analysis of NGAL and KIM-1 expression by ImageJ software. All data are presented as mean ± SD, \**p* < 0.05; \*\**p* < 0.01.

**Figure 6.**
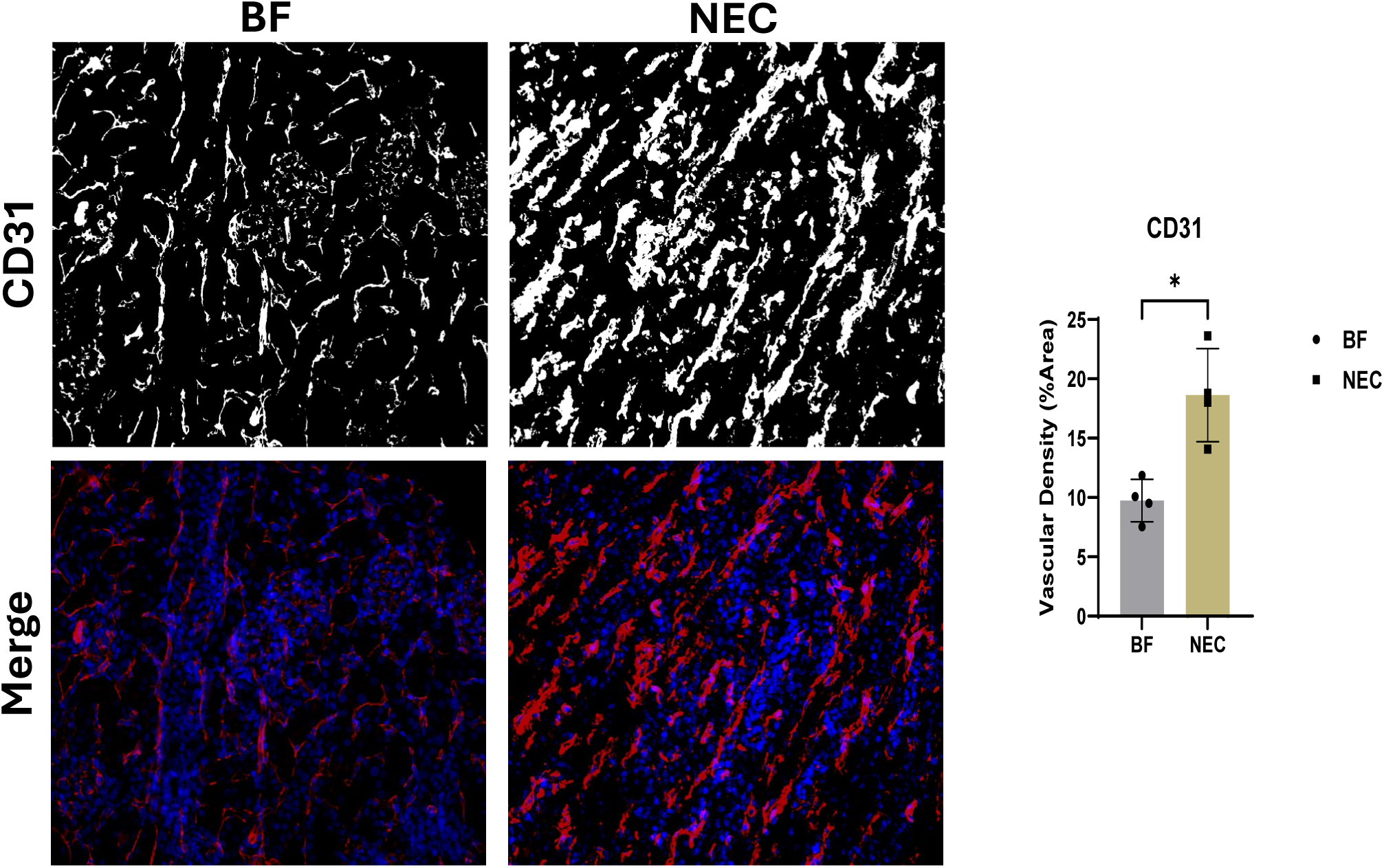
Expression of CD31 in BF and NEC kidney tissue. Representative images of immunofluorescence staining of CD31. NEC kidney sections showing strong CD31expression in comparison to BF. Quantitative analysis of CD31 expression by Image J software. All data are presented as Mean ± SD, \**p* < 0.05.

### 3.4 PAI Assessment of Tissue Oxygenation and Blood Flow in NEC-Associated AKI

PAI images of kidney tissue oxygenation and total hemoglobin were evaluated in experimental NEC and compared to BF control. Representative images for PAI assessment of kidney tissue oxygenation in 4-day old rat pups are shown in Figure 7. PAI oxygen saturation images show general reduction in the magnitude of kidney tissue oxygenation in NEC pups. Quantitative assessment of average tissue oxygenation within the kidney ROI showed 4-day old NEC pups have a significant decrease in renal oxygenation as compared to control pups (61.17% ± 1.266% versus 54.43% ± 5.573%, *p* = 0.0298), reflecting an 11% relative decrease in oxygen saturation. Representative images for PAI assessment of kidney total hemoglobin in 4-day old rat pups are shown in Figure 8. PAI images show marked increase in total hemoglobin in NEC pups. Quantitative assessment of average total hemoglobin within the kidney ROI showed that 4-day old NEC pups have a significant increase in total hemoglobin as compared to controls (24135 a.u. ± 3473 versus 44234 a.u. ± 13169, *p* = 0.0109), reflecting a 45% relative increase in total hemoglobin. These findings reflect substantial differences in renal tissue oxygenation and hemodynamics in the rat neonatal model of NEC.

**Figure 7.**
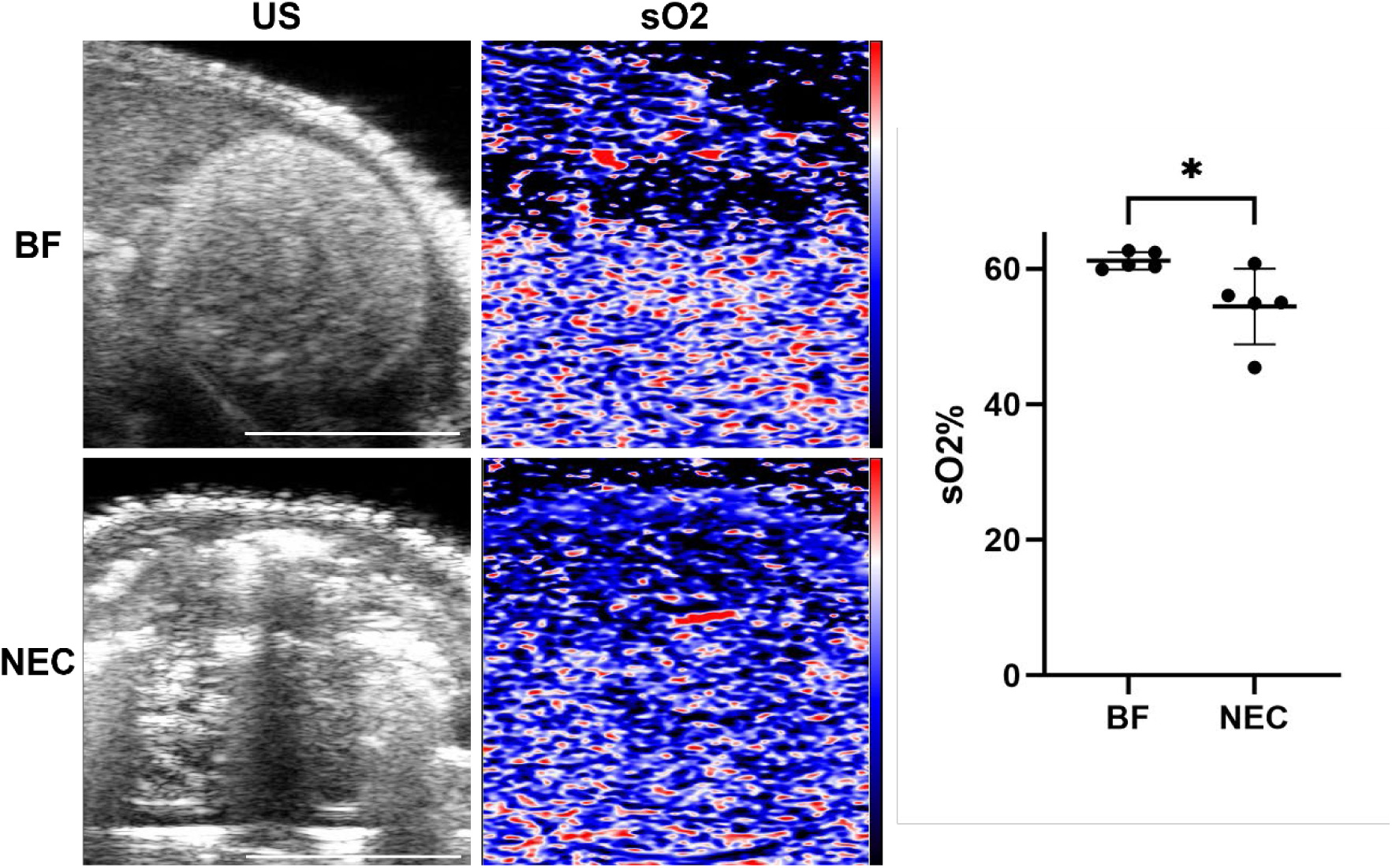
**(Left)** US and PAI images of tissue oxygenation in representative BF and NEC 4-day old rat pups. Scale bar = 5mm. **(Right)** Renal tissue oxygenation in NEC pups demonstrated a significant reduction compared to BF controls. \**p* < 0.05.

**Figure 8.**
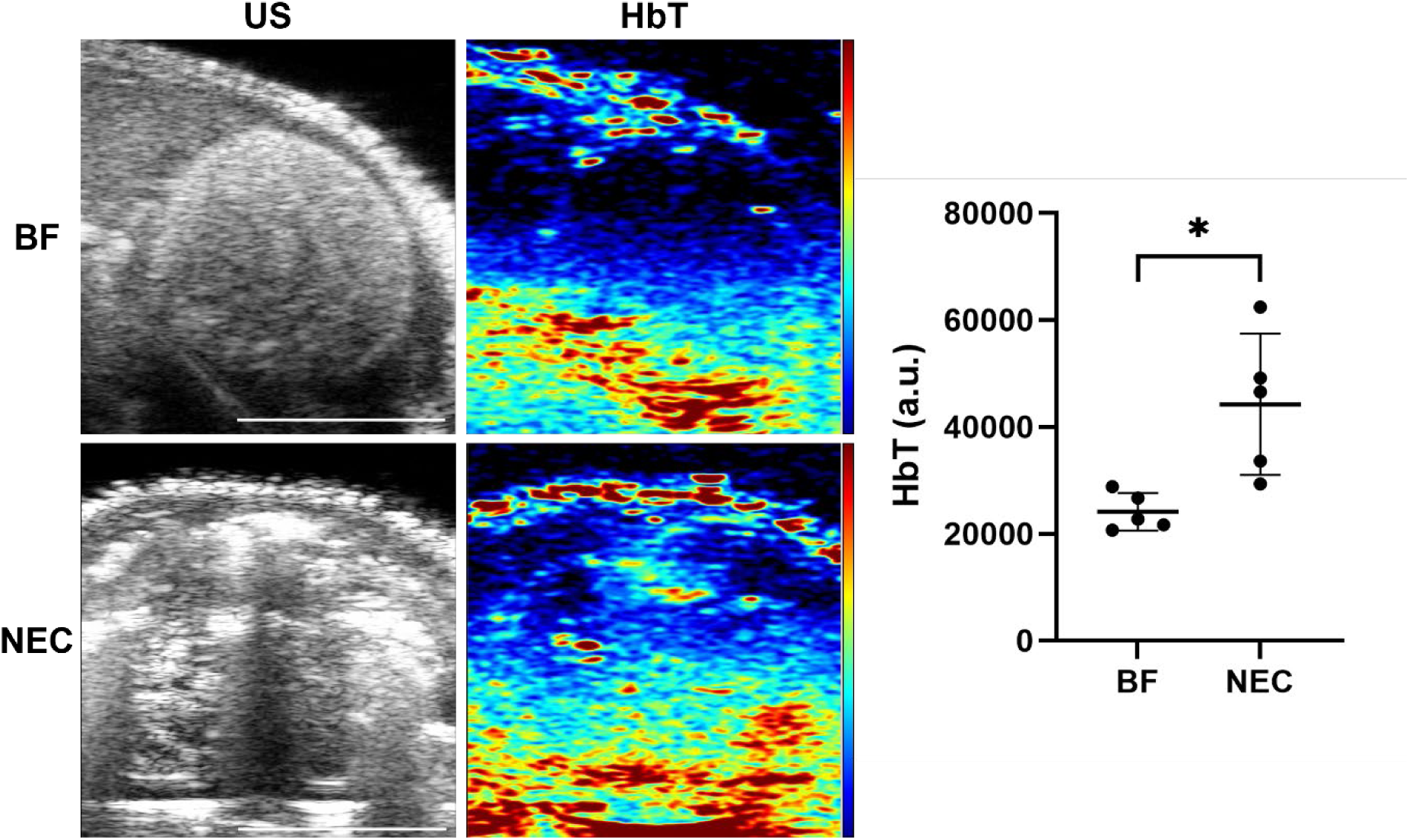
**(Left)** US and PAI images of total hemoglobin in representative BF and NEC 4-day old rat pups. Scale bar = 5 mm. **(Right)** Renal total hemoglobin in NEC pups demonstrated a significant increase compared to BF controls. \**p* < 0.05.

## 4. Discussion

NEC-associated acute kidney injury in an experimental neonatal rat model of NEC is explored in this study through an integrative multimodal approach which involves photoacoustic imaging (PAI), biochemical and molecular biomarker quantification, and histopathological analyses. PAI revealed that the NEC pups had significantly reduced oxygen saturation levels and renal tissue hypoxia as compared to BF controls. This finding is physiologically consistent with the hypoxic-ischemic mechanisms commonly attributed to NEC-associated kidney damage, which is characterized by impaired tissue perfusion and systemic microvascular dysfunction. These results are aligned with previous preclinical and clinical studies that demonstrate the incidence of AKI in NEC and its role in worsening newborn survival(10, 13, 27). Interestingly, despite the observed hypoxic changes, PAI also showed a significant increase in total hemoglobin in NEC pups as compared to BF control pups, reflecting an early acute reactive hyperemia response to renal tubular damage. This was confirmed by CD31 staining showing increased microvascular density. These findings suggest a complex hemodynamic response in our animal model of NEC-associated AKI. Similar emerging evidence in septic acute kidney injury models have suggested a role for renal hyperemia in a hyperdynamic circulation state for exacerbating early acute kidney damage(35, 36), with suggestion that dysregulation and maldistribution of intrarenal blood flow leads to localized hemodynamic/oxygenation mismatch within the kidney in early AKI(37, 38). Based on these and other findings, recent evidence suggests that multiple and sometimes overlapping mechanisms are at play in the multifactorial pathogenesis of AKI(39). However, to date studies have not been able to delineate the precise role of hemodynamic mechanisms in kidney damage due to a lack of reliable tools to simultaneously assess spatial renal blood flow and oxygenation(36). PAI offers a promising solution to this knowledge gap, with real-time noninvasive measurement of oxygenation and perfusion providing new insights into dynamic vascular response during early NEC-associated AKI.

Additionally, NEC pups showed significantly higher concentrations of inflammatory markers (TNF-α, IL-6, and IL-1β) in both plasma and renal tissue. These elevated levels confirm tubular damage and are consistent with other research demonstrating that intestinal inflammation and epithelial barrier damage induces inflammatory mediators that may lead to systemic inflammation that may exacerbate tubular injury(13). Further histopathological and immunofluorescence analyses illustrated the presence of acute tubular injury represented by tubular dilation and interstitial edema along with KIM-1 and NGAL expression in the affected tubules(21, 22, 40). CD31 staining revealed preservation of renal microvasculature architecture in BF kidneys, whereas NEC pups demonstrated elevated CD31 expression patterns indicative of endothelial dysfunction and compromised microvascular integrity. The increased CD31 positivity in peritubular and glomerular capillaries of NEC kidneys parallels the hemodynamic disturbances and microvascular dysregulation observed by PAI, suggesting that endothelial cell disruption contributes to underlying NEC-associated AKI. These findings indicate that in addition to tubular epithelial injury reflected by KIM-1 and NGAL expression, NEC causes significant endothelial dysregulation that exacerbates the multifactorial mechanisms driving AKI.

NEC-induced intestinal inflammation stimulates systemic inflammatory cascades, which consequently induce renal inflammatory pathways, resulting in structural and functional impairment, according to physiological findings derived from our molecular and immunological analyses. The primary mechanism that connects renal injury and intestinal inflammation in the gut-kidney axis is likely the activation of macrophage-derived NLRP3 inflammasome complexes, which causes IL-1 to release leading to both local and systemic inflammation(16). Furthermore, oxidative stress and microvascular dysregulation are caused by systemic inflammation, aggravating ischemia injury in renal tissue(13, 40). These associated pathophysiological mechanisms illustrate the impact of NEC on other extraintestinal organs and underline the significance of comprehensive monitoring strategies.

The adoption of photoacoustic imaging in this study offers substantial advantages compared to other conventional imaging techniques like magnetic resonance imaging (MRI), standard ultrasound, and near-infrared spectroscopy (NIRS). PAI specifically detects and quantifies tissue oxygenation and vascular perfusion by integrating optical spectral hemoglobin absorption with the spatial resolution of ultrasound and greater tissue depth than NIRS(25, 27). This functional imaging potential enables real-time and noninvasive assessments of renal oxygenation and blood flow levels in neonates to enhance our understanding of NEC-associated AKI pathophysiology with potential to improve clinical detection of AKI at an early stage(13, 27).

## Acknowledgements

We acknowledge the Preclinical Ultrasound and Photoacoustic Imaging Core of Wake Forest University School of Medicine supported in part by the Wake Forest Clinical and Translational Science Institute (NIH NCATS UL1TR001420) and the Hypertension and Vascular Research Center. We also acknowledge the Biomarker Analytical Core of Wake Forest University School of Medicine.

## Grants

These studies were supported in part by the National Institutes of Health (NIDDK K01DK125633, NIDDK R01DK135955, and NIBIB T32EB014836) and the American Gastroenterological Association Research Scholar Award in Health Disparities.

## Disclosures

The authors have no conflicts of interest to declare.

## Contributions

Conceived and designed research: AG, JW, VW; Analyzed data: AG, AS, JW, VW; Performed experiments: AG, AS, ME, NCD, JW, VW; Interpreted results of experiments: AG, AS, JW, VW; Prepared figures: AG, AS, JW; Drafted manuscript: AG; Edited and revised manuscript: All authors; Approved final version: All authors

## Data availability

The datasets generated and/or analyzed within the current study are available from the corresponding authors on reasonable request.

## Notes

### Competing Interest Statement

The authors have declared no competing interest.

### Summary of Updates

Updated histological assessments by adding more H&E images, adding CD31 immunofluorescence data, updating KIM1 and NGAL immunofluorescence data and immunofluorescence quantitative analyses.

